## Supplemental Figures for "Interpretable single-cell factor decomposition using sciRED"

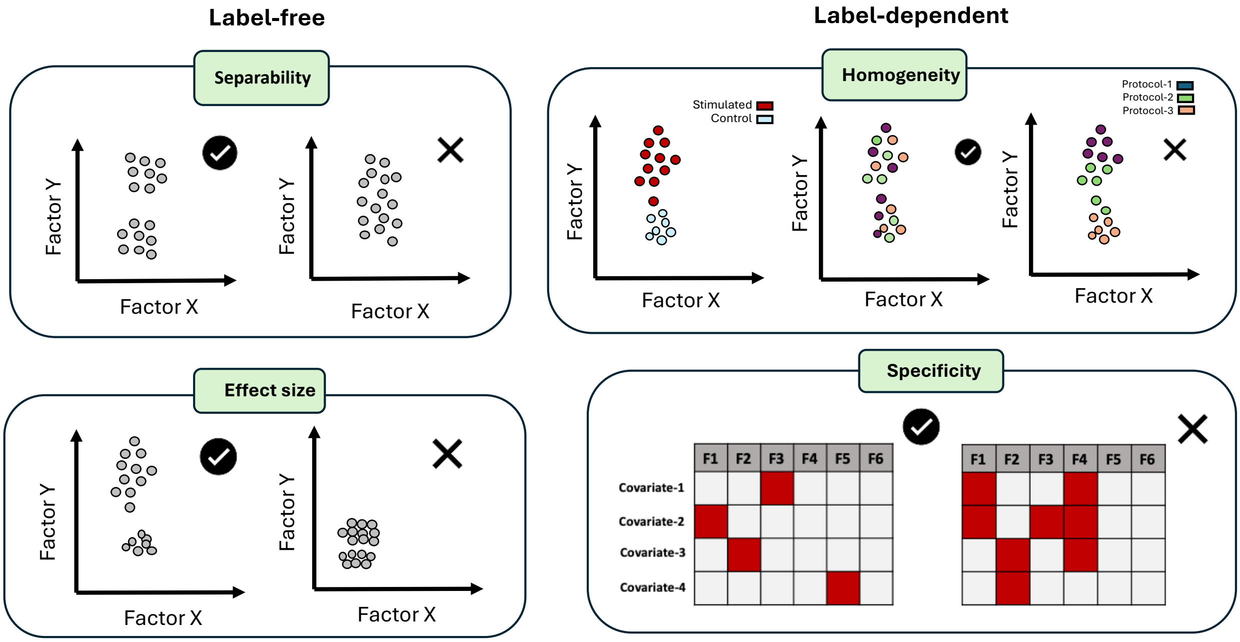


Supplementary Figure 1) Schematic overview of factor interpretability metrics

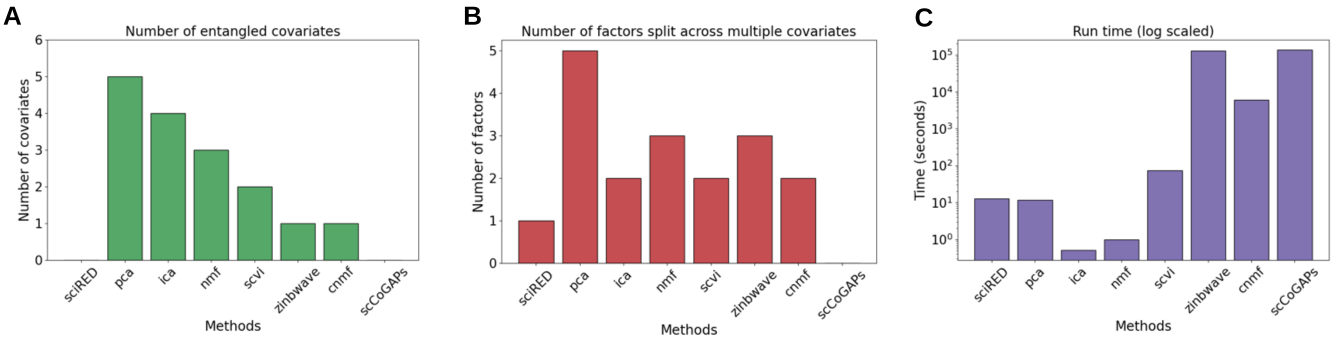


Supplementary Figure 2) Benchmarking sciRED's factor discovery on the scMixology dataset.

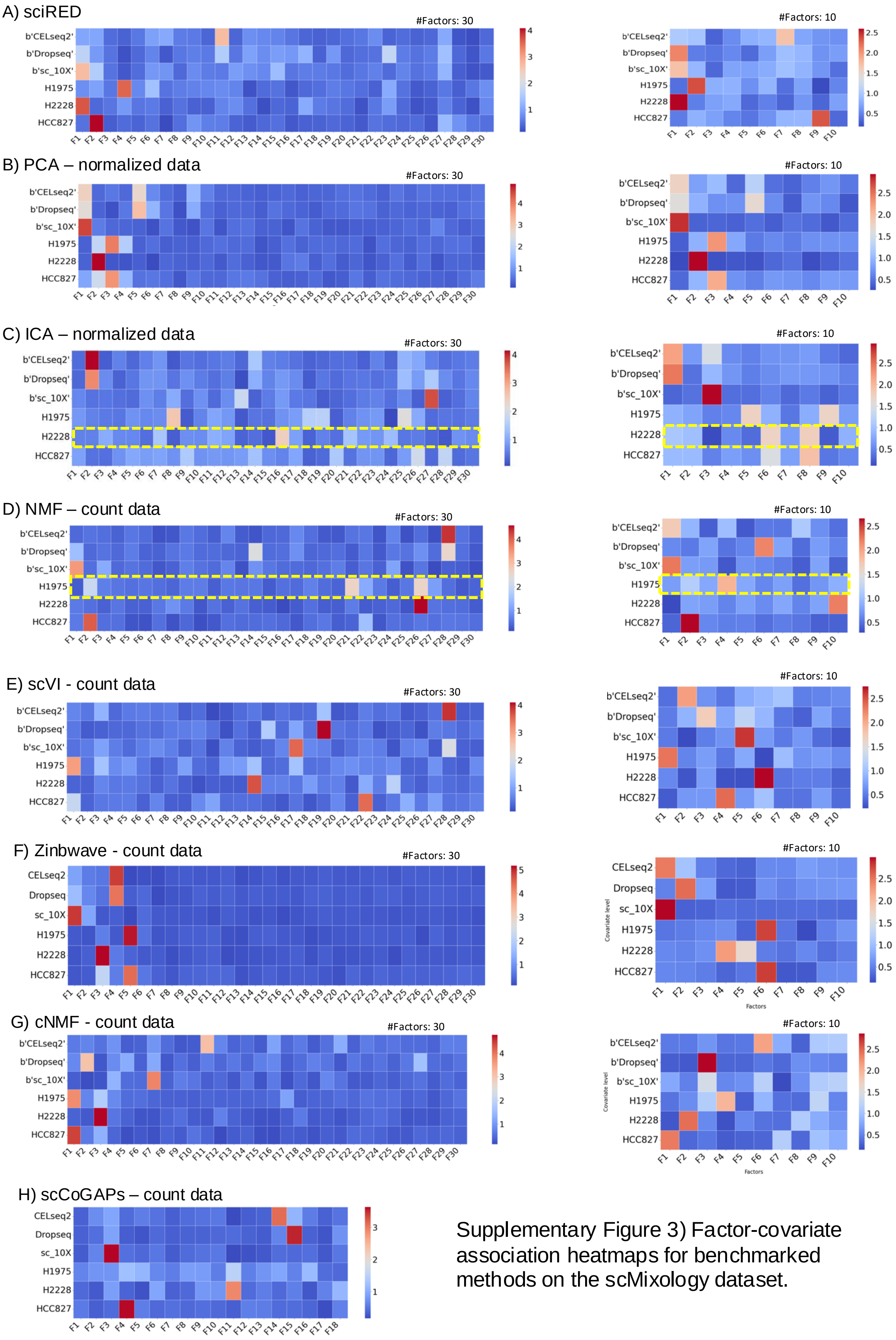


Supplementary Figure 3) Factor-covariate association heatmaps for benchmarked methods on the scMixology dataset.

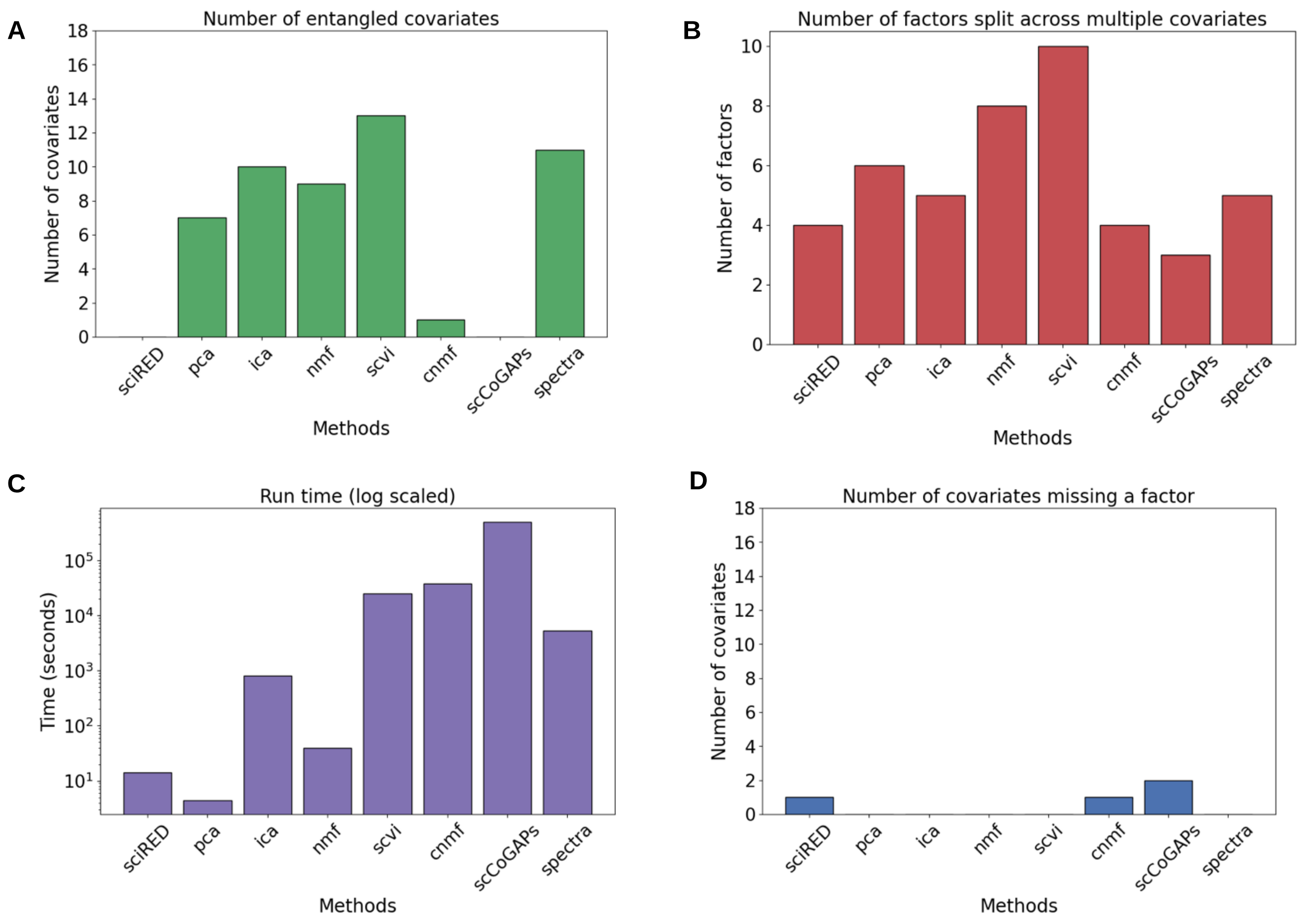


Supplementary Figure 4) Benchmarking sciRED's factor discovery step on a biological dataset.

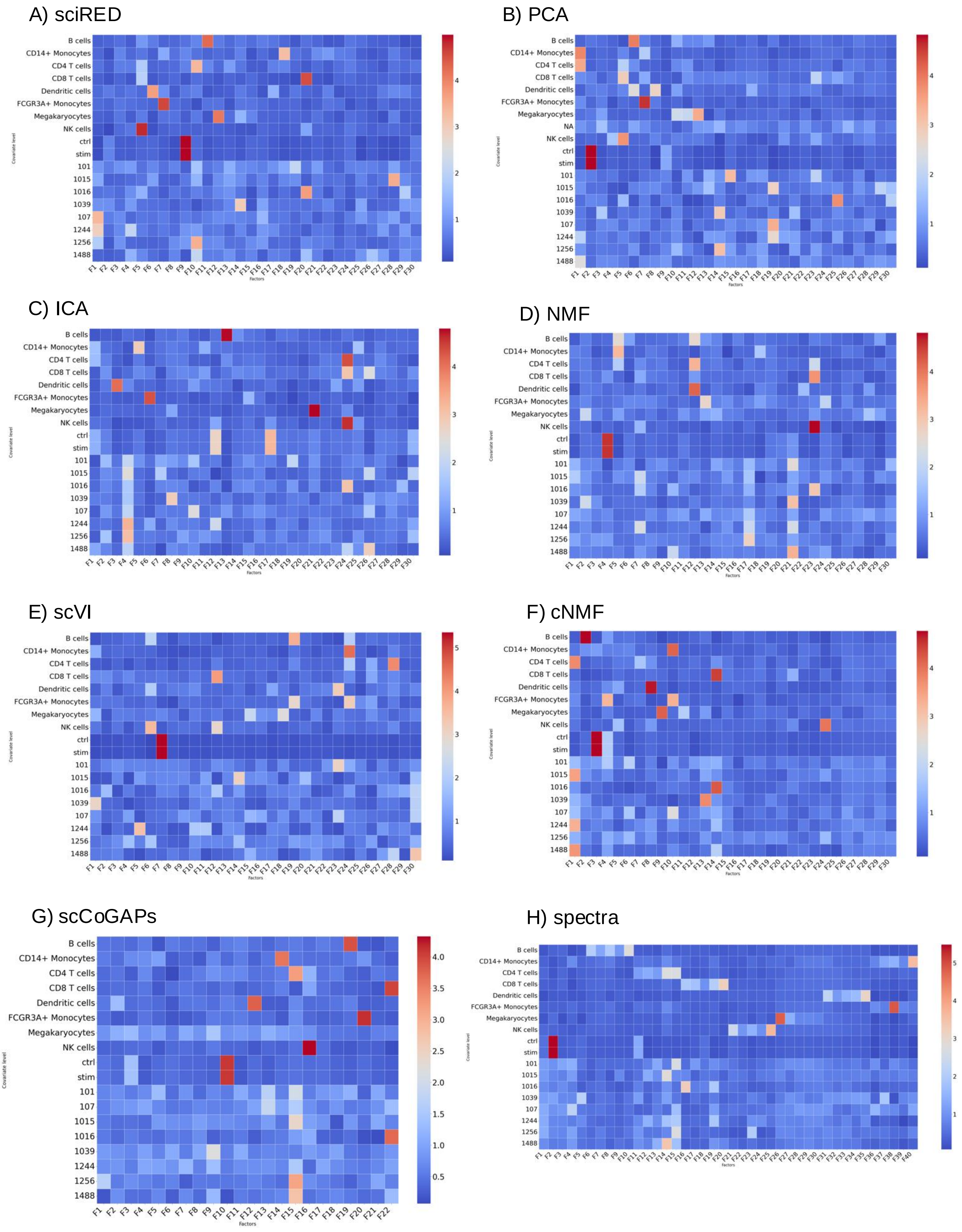


Supplementary Figure 5) Factor-covariate association heatmaps for benchmarked methods on the stimulated PBMC dataset.

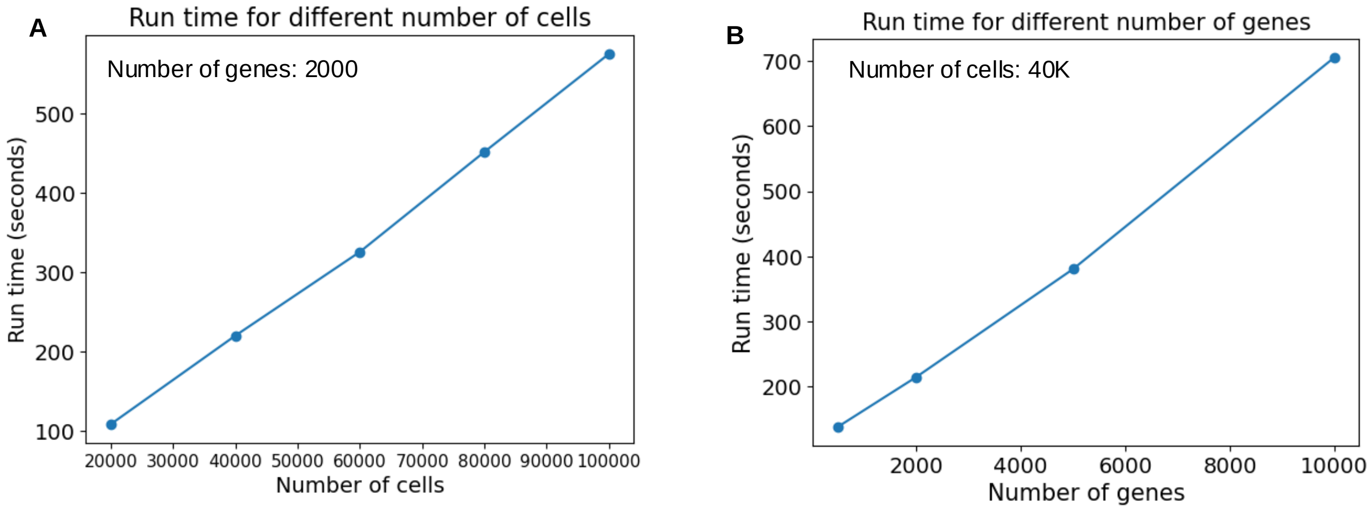


Supplementary Figure 6) Runtime analysis relative to the number of cells and genes.

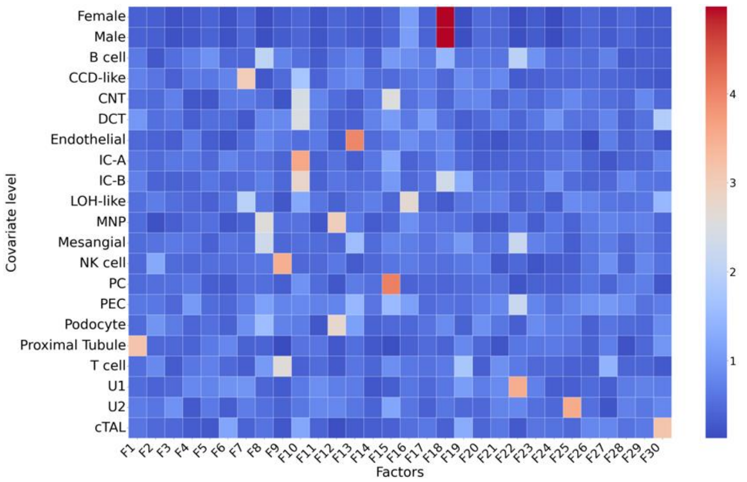


Supplementary Figure 7) The complete factor-covariate association table of the healthy kidney atlas.

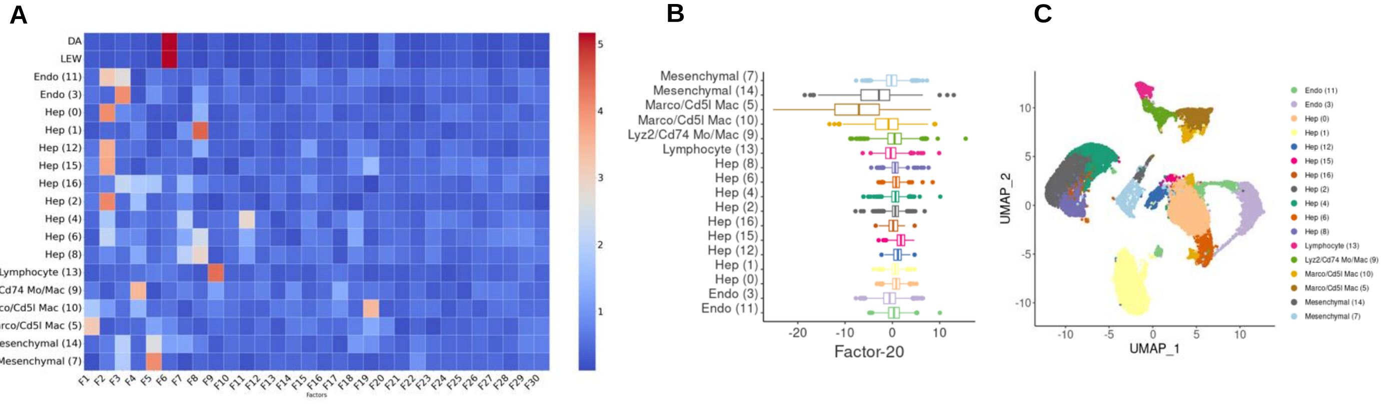


Supplementary Figure 8) Annotated healthy rat liver map

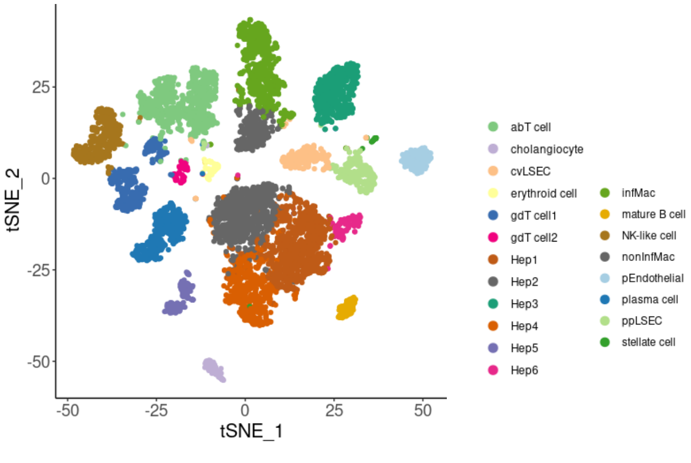


Supplementary Figure 9) Annotated healthy human liver map

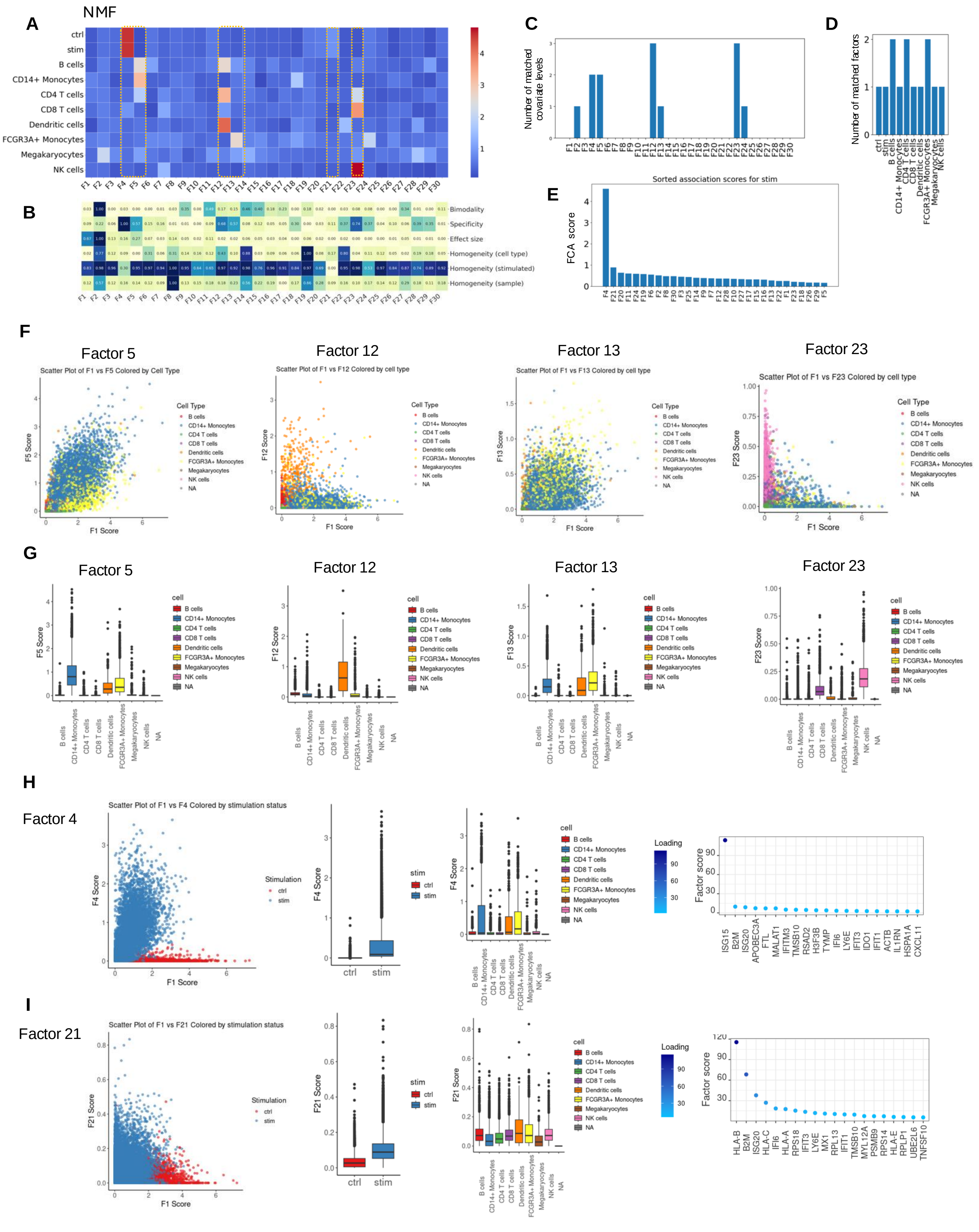


Supplementary Figure 10) Applying sciRED's interpretation and evaluation steps to alternative factor identification methods.

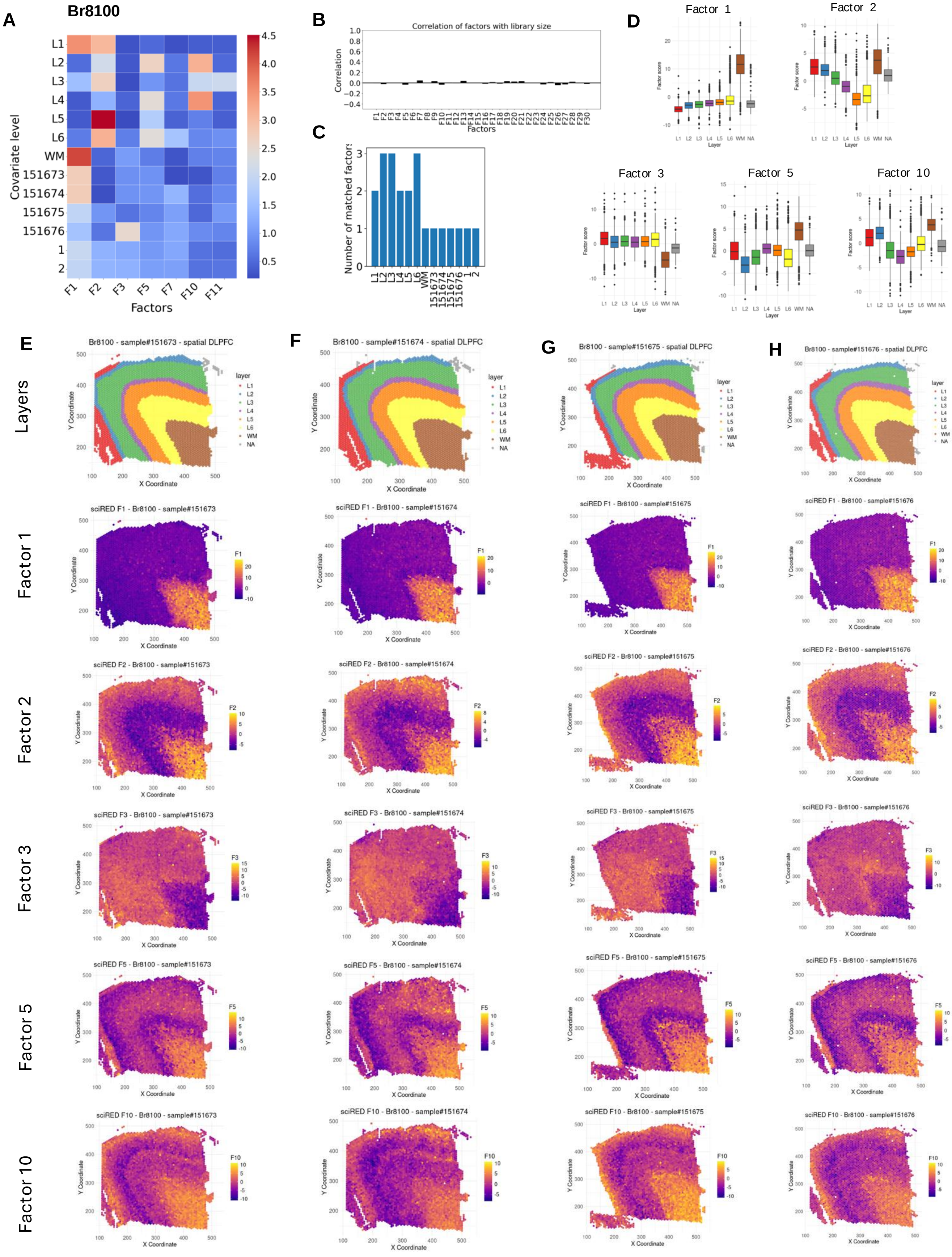


Supplementary Figure 11) Application of sciRED to spatial transcriptomics data (subject Br8100).

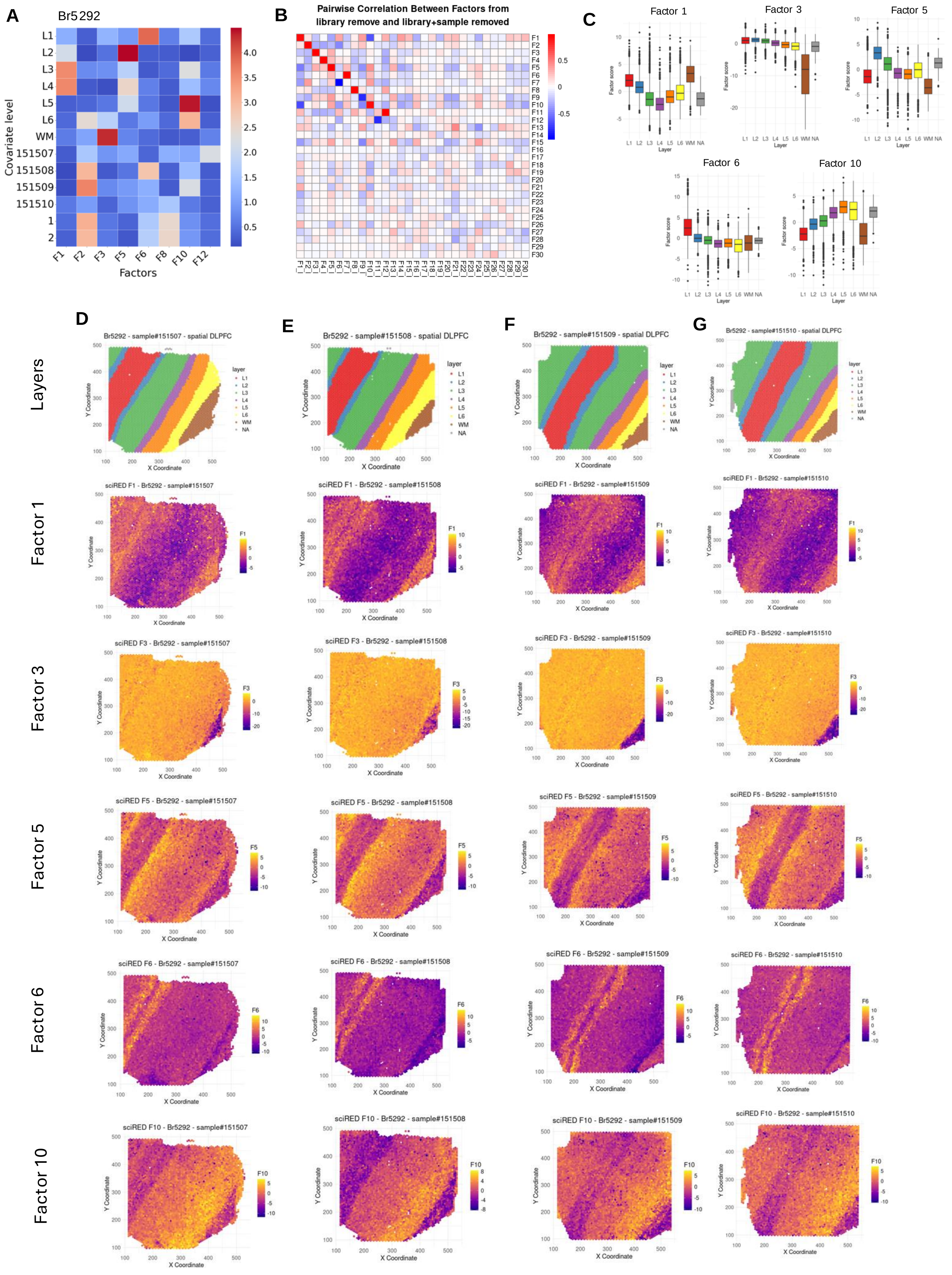


Supplementary Figure 12) Application of sciRED to spatial transcriptomics data (subject Br5292)

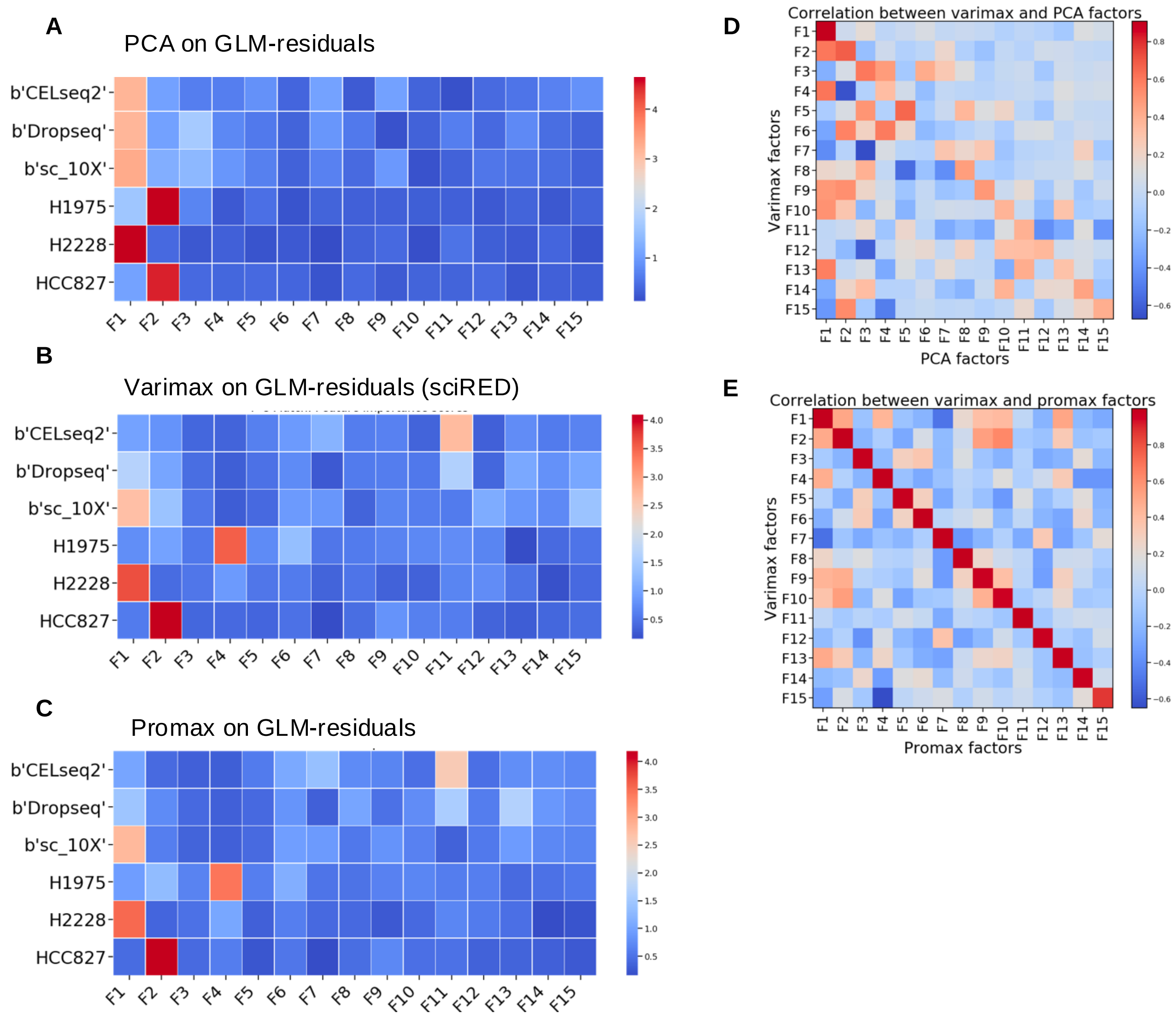


Supplementary Figure 13) Enhancement in identifying cell line-specific factors through rotations.

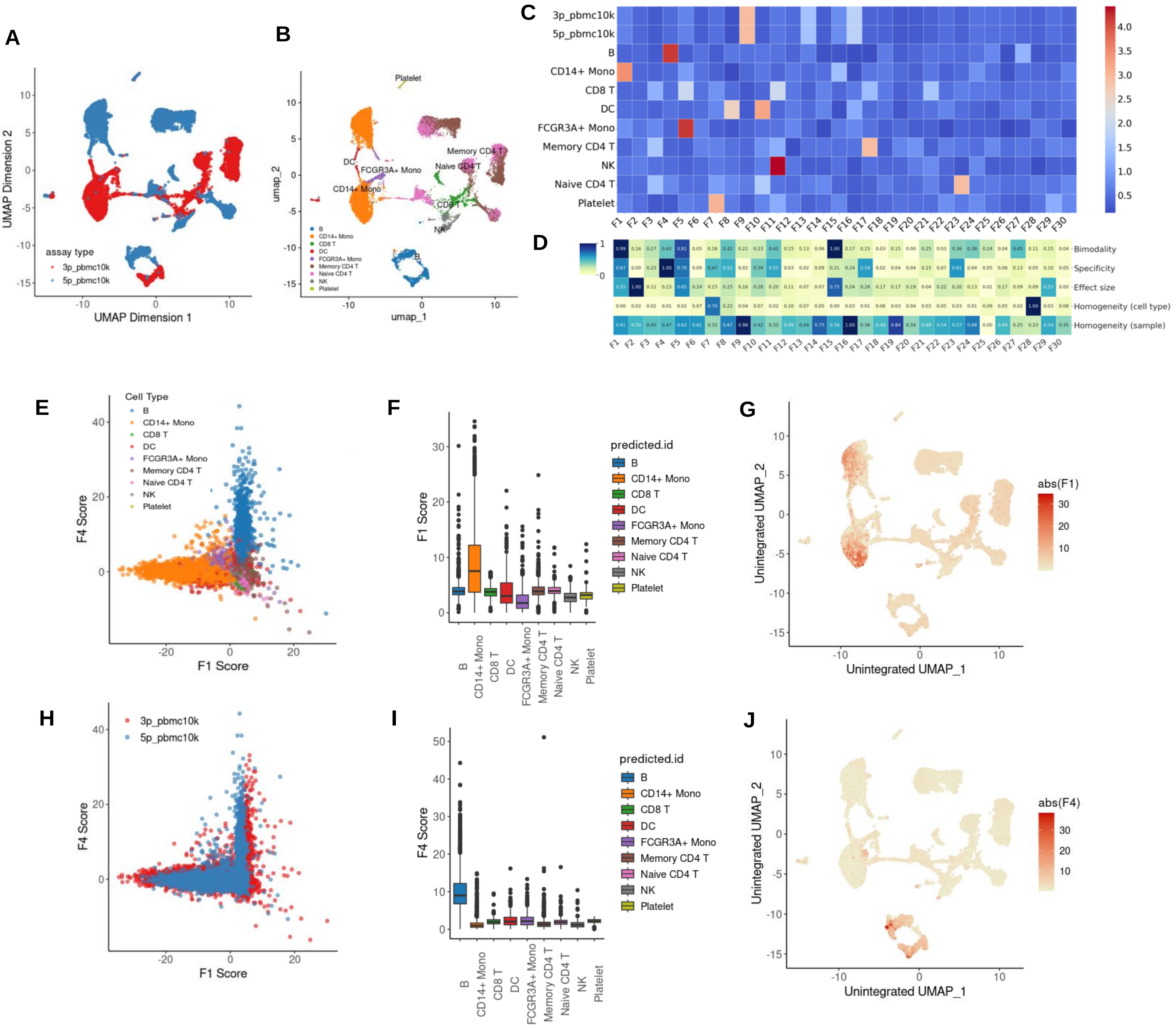


Supplementary Figure 14) sciRED performance in the presence of strong batch effects.

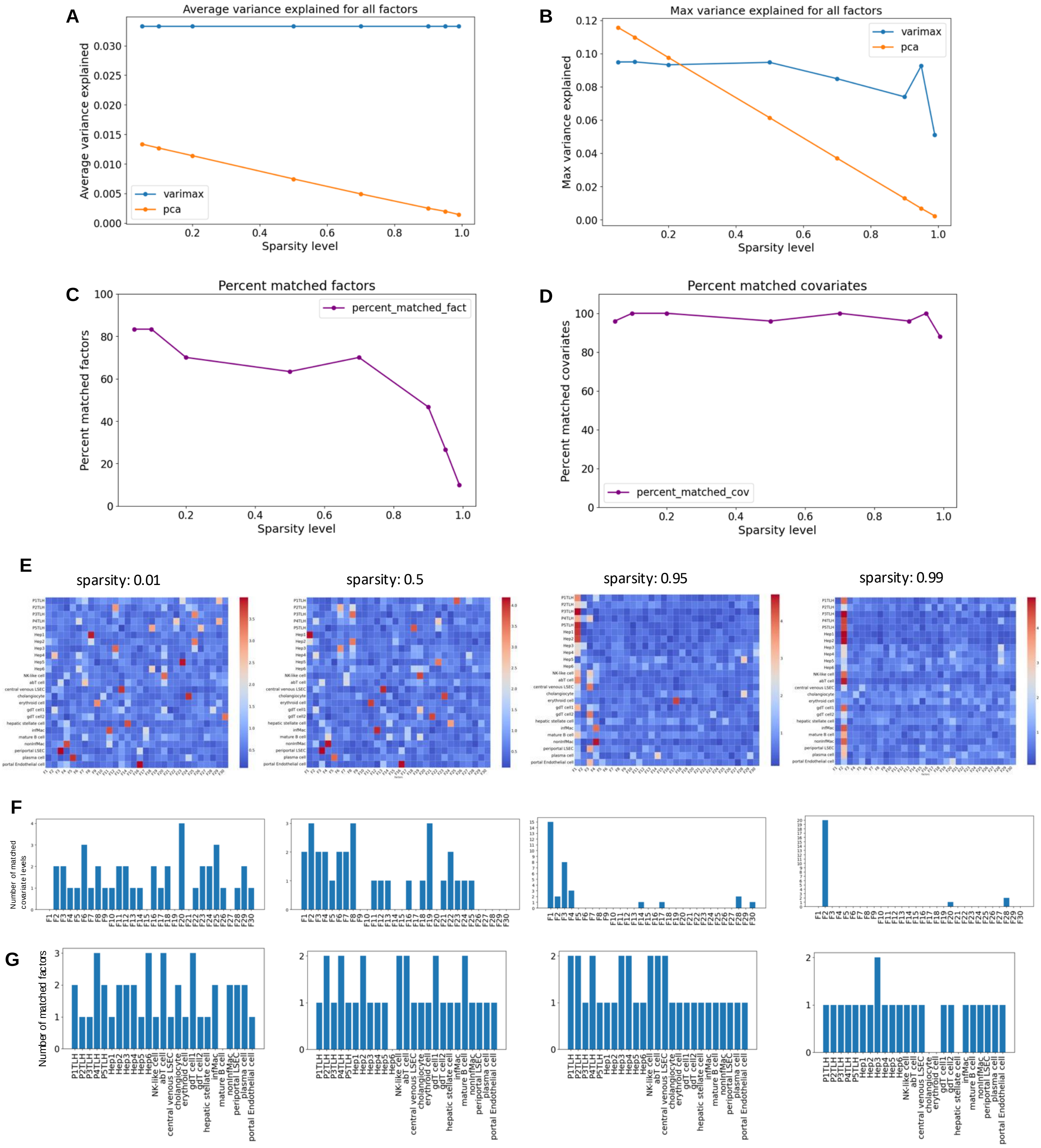


Supplementary Figure 15) Evaluation of sparsity effect on sciRED's decomposition results.

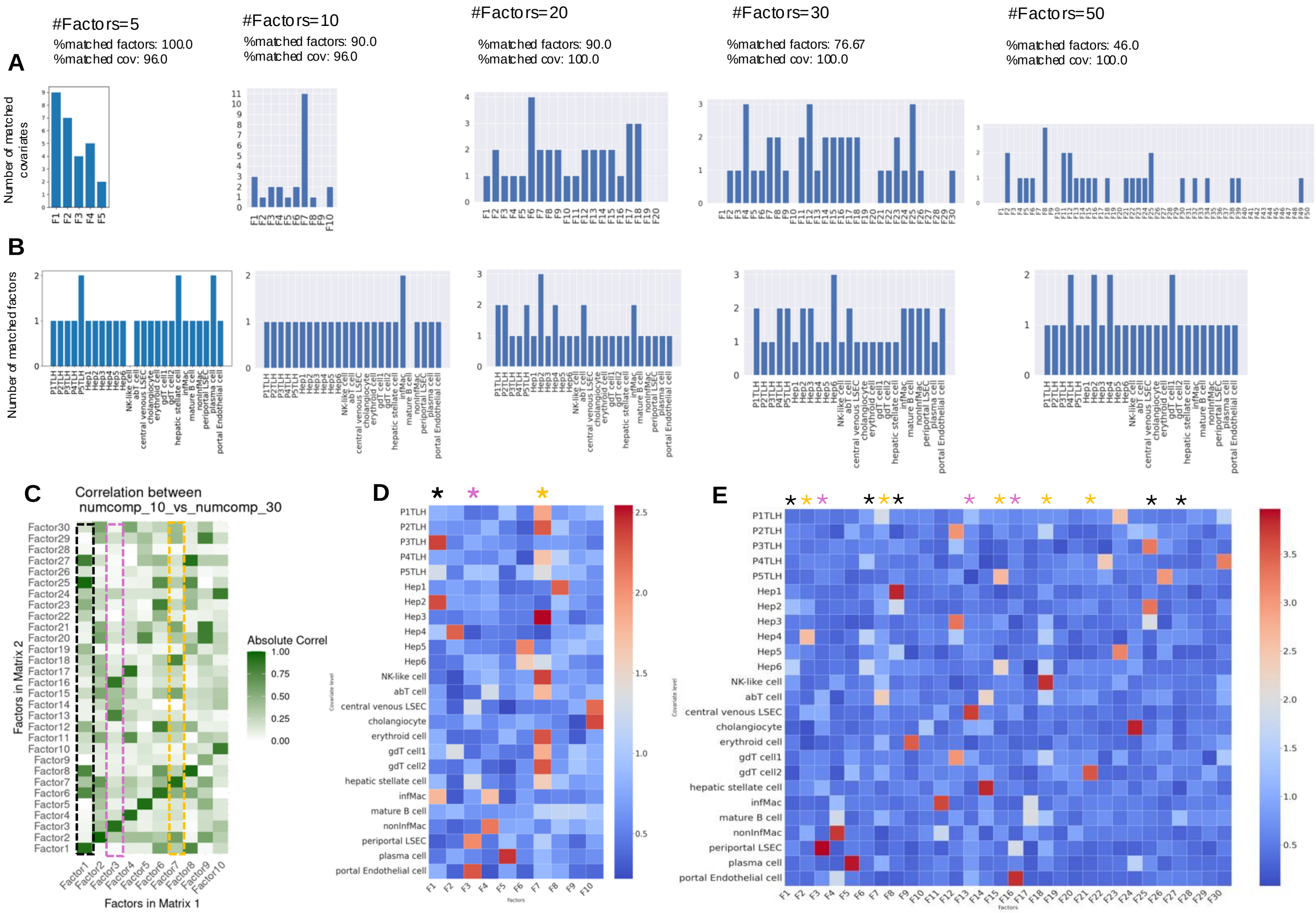


Supplementary Figure 16) Evaluating the effect of factor number on sciRED's decomposition results based on the human liver map.

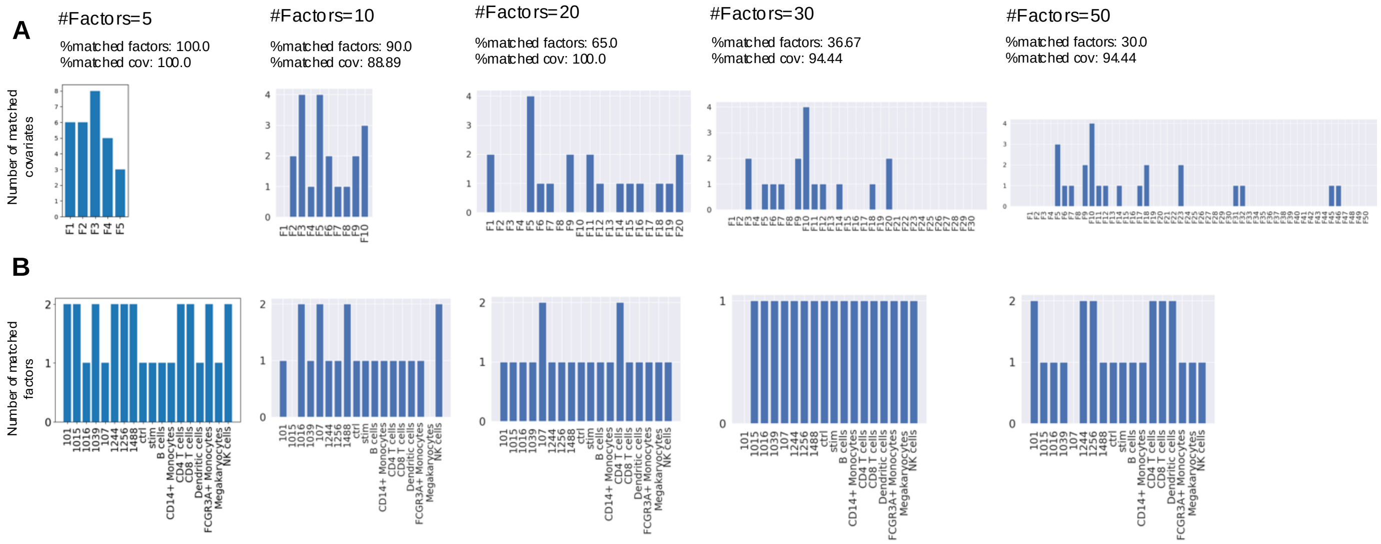


Supplementary Figure 17) Evaluating the effect of factor number on sciRED's decomposition results using the stimulated PBMC dataset.

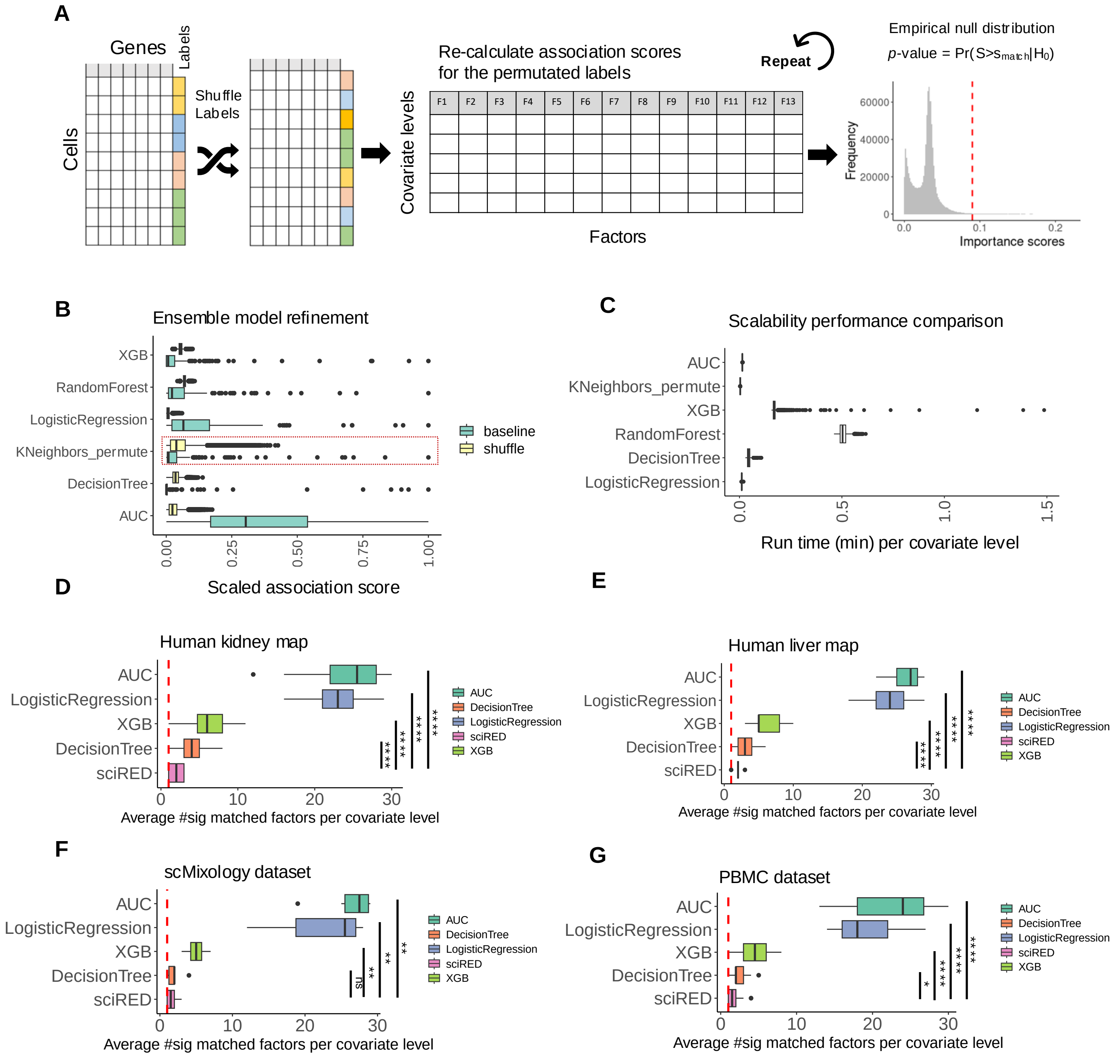


Supplementary Figure 18) Performance evaluation and benchmarking of sciRED.

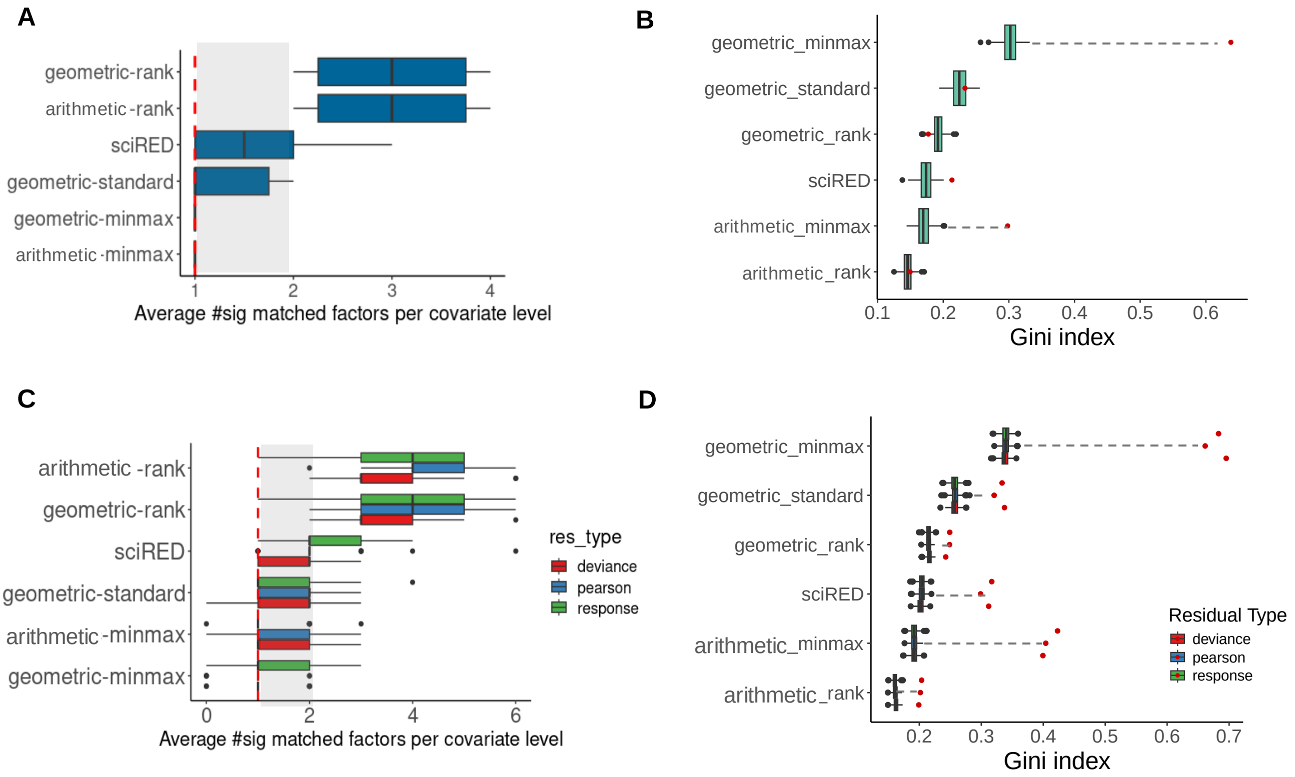


Supplementary Figure 19) Optimizing ensemble design through permutation-based comparison of scaling and mean calculation methods.

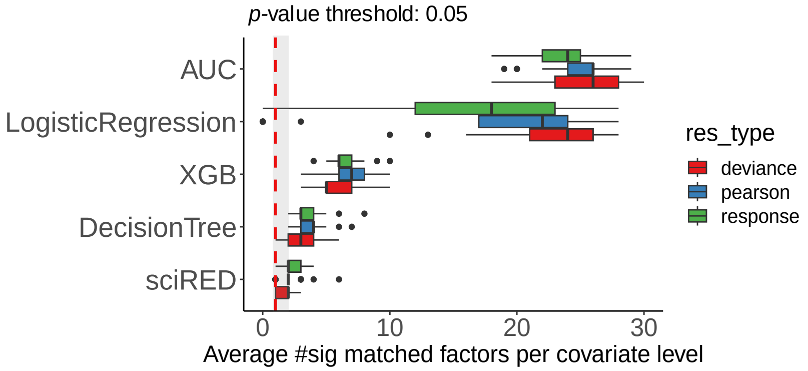


Supplementary Figure 20) Impact of residual choice on factor interpretability.

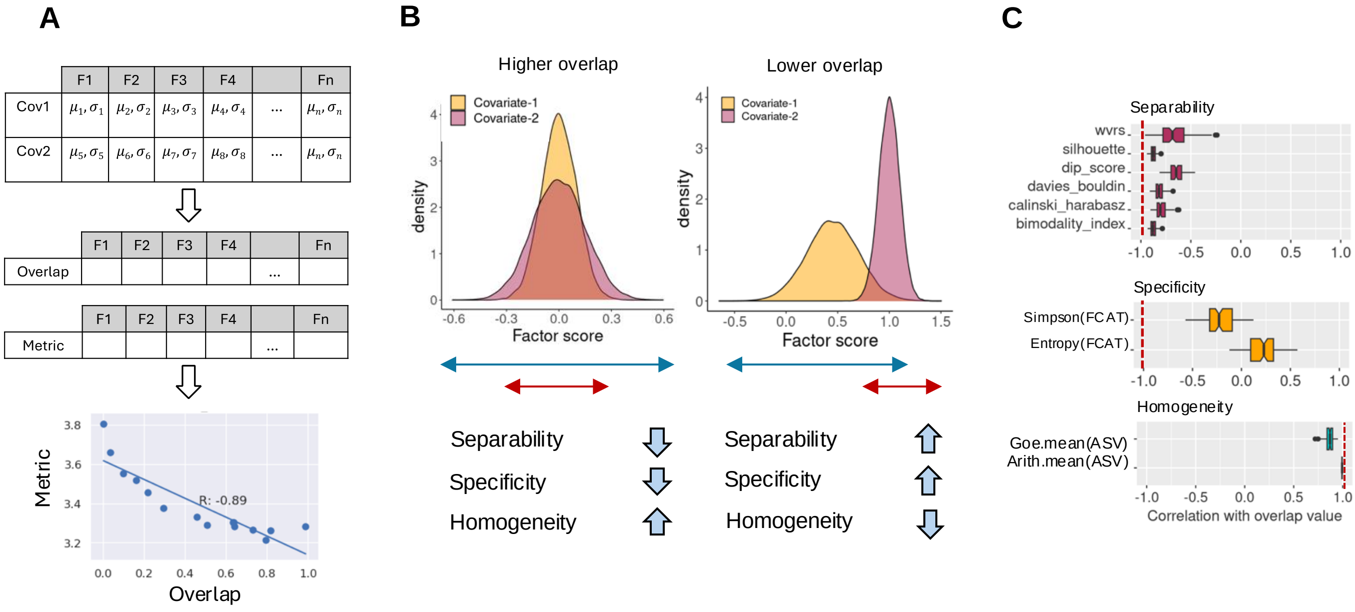


Supplementary Figure 21) Evaluating factor interpretability metrics through factor simulation
